# Control of *Staphylococcus aureus* quorum sensing by a membrane-embedded peptidase

**DOI:** 10.1101/516906

**Authors:** Chance J. Cosgriff, Chelsea R. White, Wei Ping Teoh, James P. Grayczyk, Francis Alonzo

## Abstract

Gram-positive bacteria process and release small peptides or “pheromones” that act as signals for the induction of adaptive traits including those involved in pathogenesis. One class of small signaling pheromones is the cyclic auto-inducing peptides (AIPs), which regulate expression of genes that orchestrate virulence and persistence in a range of microbes including Staphylococci, Listeria, Clostridia, and Enterococci. In a genetic screen for *Staphylococcus aureus* secreted virulence factors, we identified a *S. aureus* mutant containing an insertion in gene *SAUSA300_1984* (*mroQ*), which encodes a putative membrane-embedded metalloprotease. A Δ*mroQ* mutant exhibits impaired induction of TLR2-dependent inflammatory responses from macrophages, but elicits greater production of the inflammatory cytokine IL-1β and is attenuated in a murine skin and soft tissue infection model. The Δ*mroQ* mutant phenocopies a *S. aureus* mutant containing a deletion of the accessory gene regulatory system (Agr), wherein both strains have significantly reduced production of secreted toxins and virulence factors, but increased surface Protein A abundance. The Agr system controls virulence factor gene expression in *S. aureus* through sensing accumulation of AIP via the histidine kinase AgrC and response regulator AgrA. We provide evidence to suggest that MroQ acts within the Agr pathway to facilitate optimal processing or export of AIP for signal amplification through AgrC/A and induction of virulence factor gene expression. Mutation of MroQ active site residues significantly reduces AIP signaling and attenuates virulence. Altogether, this work identifies a new component of the Agr quorum sensing circuit that is critical for the production of *S. aureus* virulence factors.

## Introduction

Quorum sensing in Gram-positive bacteria occurs via the maturation and release of small signaling oligopeptides (1). These peptides can be linear, but may also undergo cyclic ring formation, as is known to occur with the *Staphylococcus aureus* auto-inducing peptide, AIP (2, 3). In either scenario, a precursor peptide is synthesized and processed prior to, or after export from the bacterial cell. While many signaling molecules of Gram-negative bacteria are freely diffusible, the peptides of Gram-positives generally must transit the membrane via a dedicated transporter (4, 5). After processing and transport, the peptide is either imported back into the bacterial cell or transmits signal from the extracellular environment by binding to membrane-embedded sensor histidine kinases (6). Peptide signaling culminates in a change in gene expression mediated by transcription factors that respond to the peptide. Many Gram-positive bacterial pathogens use these “quorum-sensing” peptides to induce gene expression programs that promote virulence adaptations such as competence, toxin production, biofilm formation, and establishment of persistence traits. Because of its importance in activating virulence programs in *S. aureus*, quorum-sensing inhibition has been the focus of many therapeutic initiatives (7–14).

The ability of *S. aureus* to infect host tissues and cause acute and chronic disease is partially due to its use of complex gene regulatory systems that control virulence factor gene expression (15–17). *S. aureus* employs 16 two-component systems that contribute to virulence and a range of environmental adaptations (18–25). One of the central two-component systems in *S. aureus* that regulates virulence factor production is the Accessory Gene Regulatory (Agr) system (15, 26). The components of the Agr system are encoded in an operon containing 4 open reading frames (*agrBDCA*) (3). These open reading frames encode the histidine kinase AgrC and its cognate response regulator AgrA, as well as AgrB - a protease that cleaves AgrD, the precursor peptide of AIP. The AgrD precursor peptide is ~46 amino acids in length with a central stretch of 7 to 9 amino acids that comprise mature AIP. The C-terminal charged tail of AgrD is cleaved by AgrB (27–30), which subsequently mediates peptide cyclization by inducing formation of a thiolactone linkage between the C-terminal carbonyl and the sulfur atom of a conserved cysteine side chain (29). The N-terminal 18 residues form an amphipathic α-helix that anchors the peptide to the cell membrane by lateral association and is required for processing and transport (31). The remaining processing and transport mechanisms for AIP are not fully defined, but in some *S. aureus* strains, the Type I signal peptidase SpsB is involved in peptide cleavage (32). Clinical isolates of *S. aureus* can be stratified into one of four Agr groups (I, II, III, and IV) that harbor allelic variants of AgrD, AgrB, and AgrC (3, 30, 33–35). Recognition of AIP from different Agr Types is group specific; and AIP variants (except AIP I and IV) inhibit the activity of non-cognate AgrCs (36–40). *S. aureus* strains of all Agr Types are known to cause clinical disease, therefore an understanding of divergence between Agr types is paramount to our understanding of pathogenesis and population dynamics (33, 41, 42). The final outcome of Agr activation in *S. aureus* is the production of toxins and other virulence factors that promote disease (3). Coincident with its critical role in regulating the production of virulence factors, strains of *S. aureus* with defects in Agr function are attenuated in skin and lung infection models, underscoring the significance of this regulatory system to pathogenesis (43–46).

We recently published the results of a screen for *S. aureus* virulence factors using a transposon mutant library of the methicillin resistant *S. aureus* (MRSA) strain JE2 (47). We hypothesized that identification of *S. aureus* mutants with secreted factors that positively or negatively modulate macrophage activation would yield good virulence factor candidates given the significant immunopathology associated with *S. aureus* infection and the major involvement of *S. aureus-*macrophage interactions in disease. Indeed, the screen successfully uncovered a range of previously unstudied mediators of virulence that our laboratory has been investigating 4 in recent years (47–49). This study relates to an insertion mutant identified in the above-mentioned screen, whose supernatant elicited production of high levels of IL-1β, but reduced secretion of other pro-inflammatory cytokines, including KC and IL-6, by macrophages (47). The transposon insertion disrupted a single open reading frame, *SAUSA300_1984*, which encodes for a type-II CAAX metalloprotease.

Type-II CAAX proteases are multi-pass transmembrane proteins with two conserved catalytic motifs EEXXXR and FXXXH (50). The diglutamate in the EEXXXR motif facilitates the activation of water for nucleophilic attack, whereas the histidine in the FXXXH motif is important for metal binding. Both motifs are thought to be necessary for enzyme activity. In eukaryotes, type-II CAAX proteases are involved in prenylation pathways that target proteins to the cell membrane through lipidation of a conserved cysteine within a CAAX motif (50). In bacteria, the function of CAAX proteases is more ambiguous although their activities have been linked to immunity to bacteriocins in some species (51–54). In *S. aureus*, there are at least four putative type-II CAAX metalloproteases, three of which are involved in surface protein display (SpdA, SpdB, and SpdC) (55, 56). SpdC was recently found to play a role in activation of the WalKR two-component system in *S. aureus* (57). SpdC directly interacts with the histidine kinase WalK at the division septum where it appears to have a negative effect on WalK kinase activity. SpdC interacts with WalK through its membrane-spanning domain; and due to sequence divergence from other type-II CAAX proteases in *S. aureus* its function likely does not require proteolytic activity (55, 57). Additional bacterial two-hybrid studies suggest SpdC interacts with *S. aureus* histidine kinases SaeS, SrrB, VraS, GraS, and ArlS, but not the histidine kinase of the Agr system, AgrC. Whereas SpdA-C orchestrates cell wall homeostasis in a manner that is independent of catalytic activity, the fourth type-II CAAX protease, 1984, appears to be functionally distinct and does not modulate cell wall integrity in any known way (55). To date, no additional functions for 1984 have been described. Though not yet investigated in great detail, at least one other study has implicated a type-II CAAX protease in the regulation of virulence gene expression. The CAAX protease of Group B *Streptococcus*, Abx1, acts as a regulator of the CovSR two-component system through its histidine kinase CovS, where overexpression of Abx1 promotes activation of CovS (58). Thus, there is mounting evidence for a conserved function of type-II CAAX proteases in the regulation of two-component system signaling in bacteria, which in these two instances appears to operate through interaction with a histidine kinase and is independent of catalytic activity.

In this work, we report the identification of SAUSA300_1984, named <u>M</u>embrane Protease <u>R</u>egulator <u>o</u>f Agr <u>Q</u>uorum Sensing, MroQ. We determine that MroQ promotes TLR2-mediated activation of innate immune cells while preventing induction of IL-1β by primary murine macrophages in vitro. This altered inflammatory profile is recapitulated in in vivo infection models where MroQ promotes inflammatory pathology, cytokine secretion, and bacterial persistence in a model of skin and soft tissue infection. Further investigation into the mode of action of MroQ determined that it acts to regulate the activity of the Agr quorum sensing system as evidenced by reduced secreted virulence factors and increased surface protein expression in a Δ*mroQ* mutant. Our data suggest that MroQ may interface with the Agr system at the level of peptide processing or transport to blunt activation of the AgrC/A two-component system, which remains otherwise responsive in a Δ*mroQ* mutant background. Altogether, our data support a model whereby MroQ controls gene expression at the level of the Agr system, with significant impacts on the virulence potential of *S. aureus.*

## Results

### MroQ is a predicted type-II CAAX protease that blunts TLR2-mediated recognition of *S. aureus* by macrophages

The *mroQ* open reading frame is located within the core genome of *S. aureus* and is positioned four open reading frames upstream of the Agr system (Fig 1A). The gene is flanked by *sdrH* and *groEL/ES*, which encode the serine aspartate repeat family protein SdrH (59) and the GroEL/ES stress response chaperone system (60) respectively (Fig 1A). Phyre2 prediction software analysis of the MroQ amino acid sequence suggest a topology containing seven transmembrane helices with the N-terminus positioned in the cytosol and C-terminus in the extracellular space, though the weaker predictive strength of the topology analysis at the N and C-terminus may suggest alternative arrangements in the membrane (Fig 1B). Consistent with its annotation as a type-II CAAX protease, MroQ contains a conserved diglutamate motif EEXXXR (E141-E142) and FXXXH (H180) metal coordination motif that shares sequence identity with previously characterized CAAX proteases SpdA and SpdB, with weaker conservation compared to SpdC, which is unlikely to have catalytic activity (Fig 1C). We previously determined that supernatant derived from a *SAUSA300_1984* transposon insertion mutant compromises macrophage activation (47). To begin interrogating the function of MroQ in greater detail, we first generated an in-frame deletion mutant and complementation strain that expresses *mroQ* from an ectopic site in the *S. aureus* chromosome driven by its predicted native promoter (61). WT, Δ*mroQ,* and Δ*mroQ* + *mroQ* strains replicate identically in tryptic soy broth (TSB) and Roswell Park Memorial Institute (RPMI) medium (Fig S1A-B). We found that similar to the transposon insertion mutant, supernatant derived from a Δ*mroQ* mutant enhanced production of IL-1β by murine bone marrow macrophages (BMM), but reduced production of the inflammatory cytokine, IL-6, and the neutrophil chemotactic factor, KC (Fig 1D). The perturbations in IL-6 and KC production were dependent on Toll-like receptor (TLR) mediated recognition of *S. aureus*, as evidenced by elimination of IL-6 and KC production in MyD88^−/−^ and TLR2^−/−^ macrophages, but not TLR4^−/−^ macrophages (Fig 1D and S2A-B). In contrast, the enhanced IL-1β production did not depend on TLRs (Fig 1D and S2A-B). These data indicate MroQ bears similarities to type-II CAAX proteases, is positioned in close proximity to the *agr* locus in the *S. aureus* chromosome, and perturbs the activation of innate immune cells in culture through both TLR-dependent and independent mechanisms.

**Figure 1.**
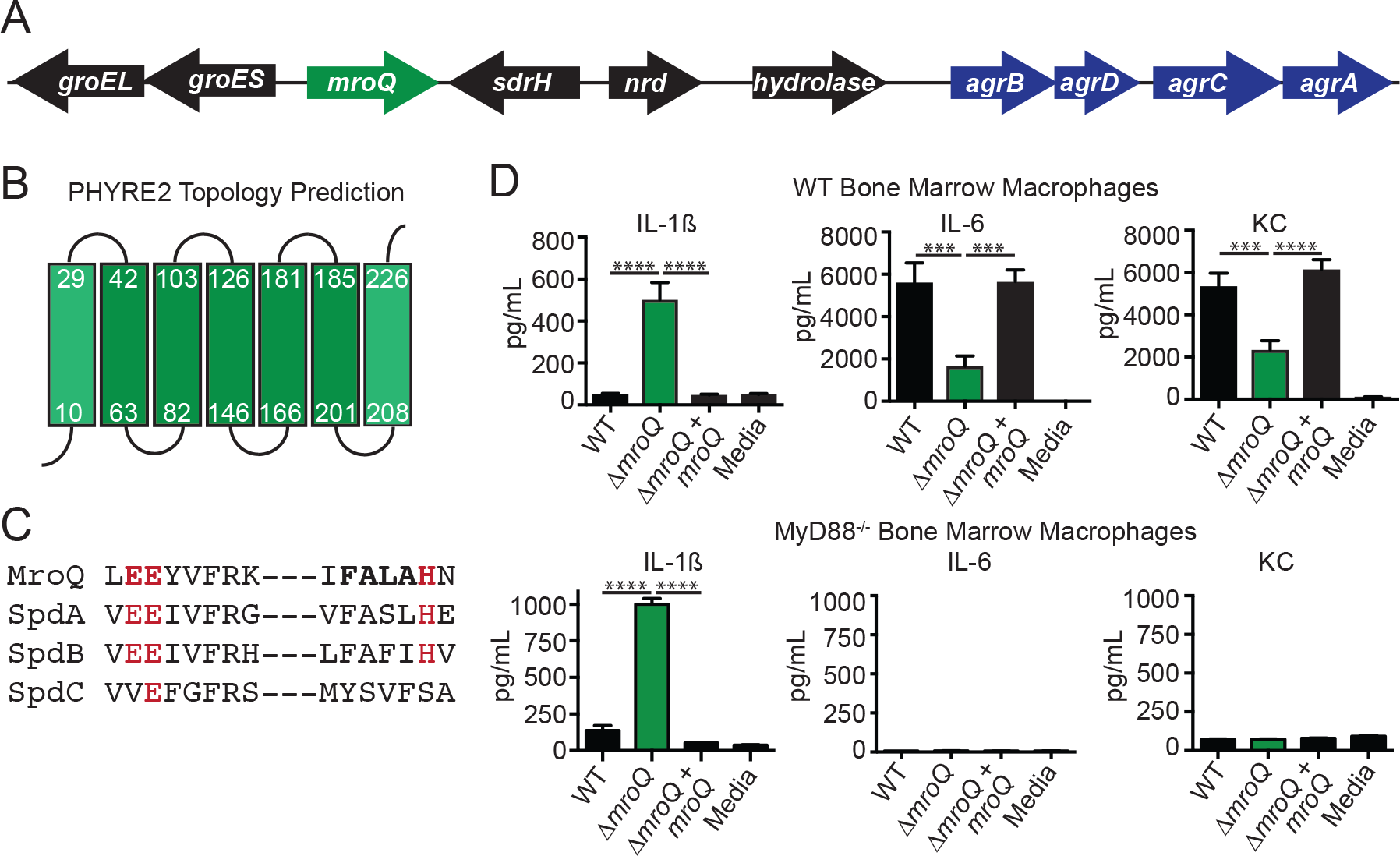
MroQ is a putative membrane-embedded protease that affects macrophage activation by *S. aureus* secreted factors. (A) Illustration of the gene arrangement surrounding *mroQ* and its proximity to the *agr* operon. *nrd* – nitroreductase family protein; *sdrH* serine aspartate repeat family protein; *hydrolase* – carbon-nitrogen family hydrolase. (B) Phyre2 topology prediction analysis of MroQ within the membrane, Green boxes – predicted α-helices. (C) Amino acid sequence alignment of the EEXXXR and FXXXH motifs that comprise the Type-II CAAX protease active site among four predicted Abi proteins of *S. aureus*. (D) IL-1β, IL-6, and KC production (pg/mL) by BMM after addition of supernatant from WT, Δ*mroQ*, or Δ*mroQ*+*mroQ* grown in TSB. Top panels, WT BMM; bottom panels, MyD88^−/−^ BMM. Data shown are from one of at least three experiments conducted in triplicate. Means ± SD are shown (n = 3). ***, *p*<0.001; ****, *p*<0.0001 by 1-way ANOVA with Tukey’s post-test.

**Figure 2.**
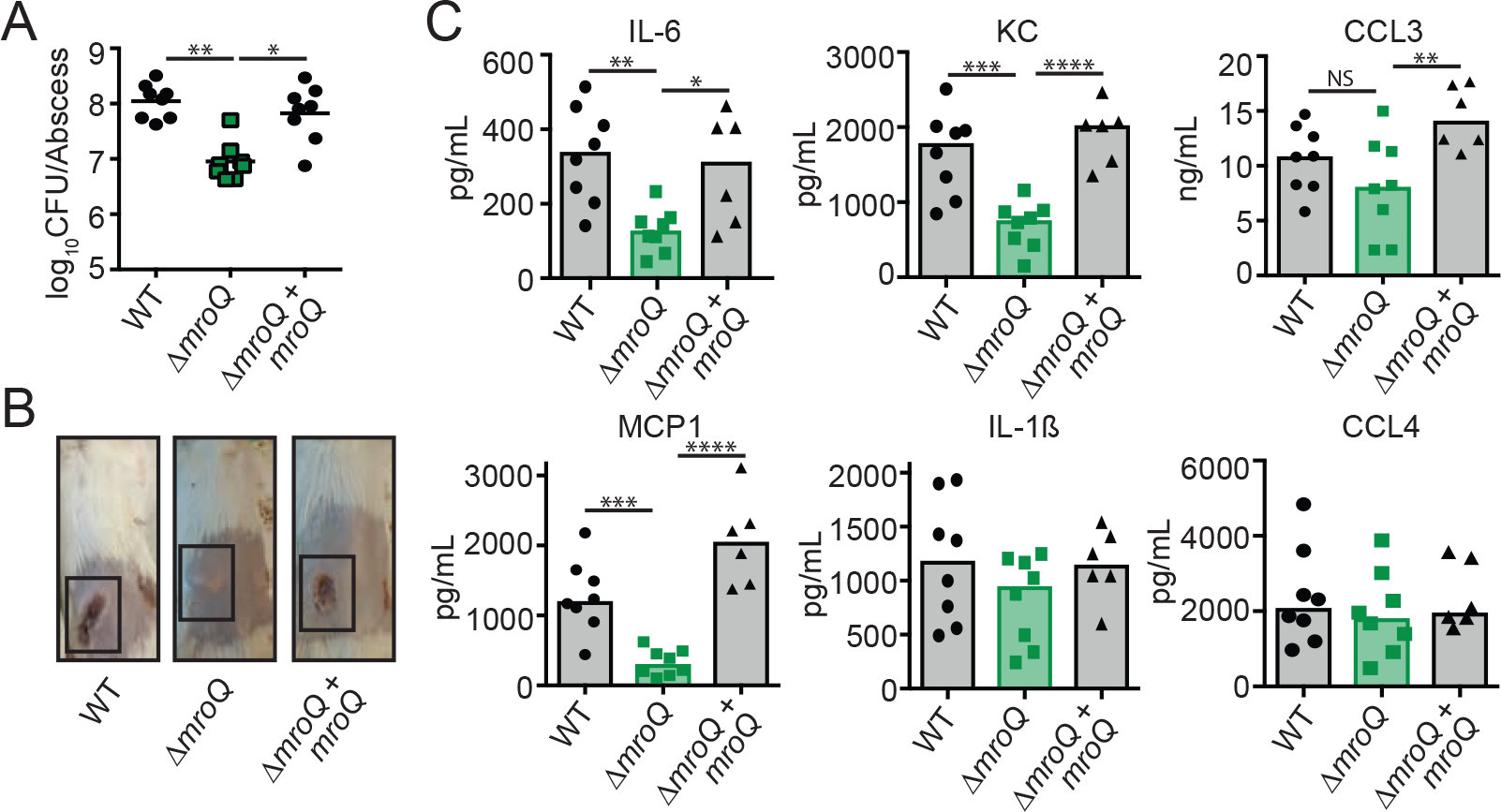
MroQ is important for *S. aureus* skin and soft tissue infection. (A) Bacterial burden in skin abscesses of mice 120 hours post-infection with WT (n=8), Δ*mroQ* (n=8), Δ*mroQ*+*mroQ* (n=8). *, *p*<0.05; **, *p*<0.01; by non-parametric 1-way ANOVA (Kruskal-Wallis Test) with Dunn’s post-test. (B) Representative images of skin abscesses 120 hours post-infection with WT, Δ*mroQ*, and Δ*mroQ*+*mroQ*. (C) IL-6, KC, MCP1, IL-1β, CCL3, and CCL4 levels in abscess homogenates of mice infected with WT (n=8), Δ*mroQ* (n=8), Δ*mroQ*+*mroQ* (n=6). *, *p*<0.05; **, *p*<0.01; ***, *p*<0.001, and ****, *p*<0.0001; by non-parametric 1-way ANOVA (Kruskal-Wallis Test) with Dunn’s post-test.

### MroQ is required for *S. aureus* virulence

Given the dramatic effect of the Δ*mroQ* mutant on macrophage activation and the predicted effects this might have on the inflammatory milieu during infection, we reasoned that the mutant would be compromised for virulence. To test this possibility, we infected mice intradermally with WT, Δ*mroQ*, and Δ*mroQ* + *mroQ* strains. There were no differences in bacterial CFU between strains at 72 hours post-infection (Fig S3A). However, significant dermonecrosis was evident in mice infected with WT and Δ*mroQ* + *mroQ* strains, but not those infected with the Δ*mroQ* mutant (Fig S3B). Consistent with reduced inflammatory tissue damage, mice infected with a Δ*mroQ* mutant showed significantly reduced levels of the inflammatory chemokine MCP-1 and trends toward reduced levels of IL-6, KC, CCL3, and CCL4 at 72 hours (Fig S3C). At 120 hours post-infection, 10-fold fewer CFU were recovered from the abscesses of mice infected with a Δ*mroQ* mutant (Fig 2A). Gross pathological signs of disease including large, ruptured, dermonecrotic abscesses were not observed in Δ*mroQ* mutant-infected mice and the abscesses remained closed with little evidence of inflammation (Fig 2B). There were significant reductions in IL-6, KC, MCP1, and CCL3 at the abscess site, consistent with in vitro studies using BMM (Fig 1D and 2C). IL-1β levels were equally elevated in all infection conditions compared to 72 hours (Fig 2C and S3C). These data indicate that MroQ is important for *S. aureus* skin and soft tissue infection.

### A Δ*mroQ* mutant phenocopies a Δ*agr* mutant

*mroQ* was identified in a screen that used secreted factors to probe innate immune cell activation (47). Therefore, we sought to test if a Δ*mroQ* mutant had notable differences in its secretome that might explain BMM activation defects and reduced virulence. Exoprotein profile analysis indicated the levels of secreted proteins are reduced in Δ*mroQ* mutant supernatants (Fig 3A). This defect in protein secretion closely resembled the profiles of a *S. aureus* strain with a deletion of the *agr* locus, Δ*agr* (Fig 3A). Exoprotein analysis of a Δ*mroQ* Δ*agr* double mutant revealed identical profiles to that of either a Δ*mroQ* or Δ*agr* deletion (Fig 3A). Furthermore, Δ*mroQ*, Δ*agr*, and Δ*mroQ* Δ*agr* supernatants induced BMM production of IL-1β, but led to reduced levels of IL-6 and KC with the same observed dependency on TLR2 (Fig 3B and S4A-C). Incidentally, both *mroQ* and *agrB* transposon insertion mutants were identified as inducers of IL-1β secretion in our prior transposon mutant library screen (47).

**Figure 3.**
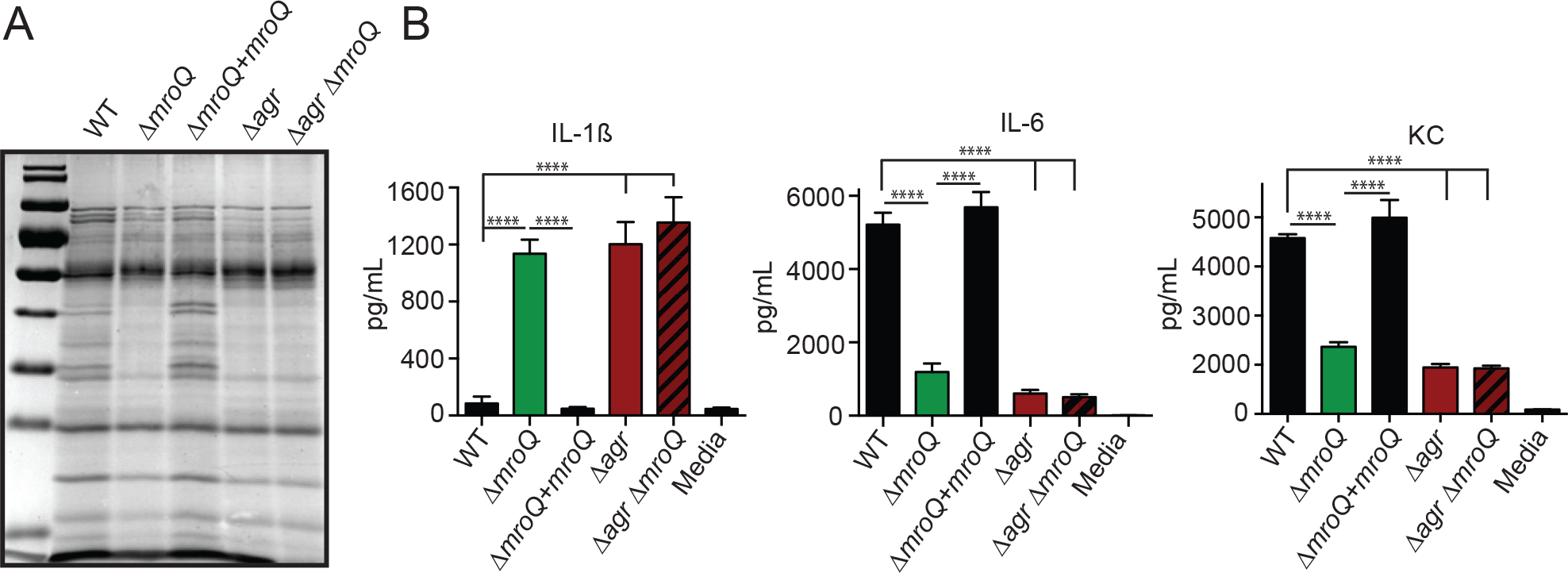
A Δ *mroQ* mutant resembles a Δ *agr* mutant for protein secretion and perturbations in macrophage activation. (A) TCA-precipitated exoproteins from WT, Δ*mroQ*, Δ*mroQ*+*mroQ*, Δ*agr*, and Δ*agr* Δ*mroQ* strains collected after growth in TSB. (B) IL-1β, IL-6, and KC production (pg/mL) by BMM after addition of supernatant from WT, Δ*mroQ*, Δ*mroQ*+*mroQ*, Δ*agr*, and Δ*agr* Δ*mroQ* grown in TSB. Data shown are from one of at least three experiments conducted in triplicate. Means ± SD are shown (n = 3). ****, *p*<0.0001 by 1-way ANOVA with Tukey’s post-test.

Given the similarities between Δ*mroQ* and Δ*agr* mutants for defects in protein secretion and secreted factor-mediated perturbations in macrophage activation, we sought to determine whether or not a Δ*mroQ* mutant exhibits other common phenotypes associated with a Δ*agr* mutant. The Agr system regulates toxin secretion and surface protein production (3). Therefore, we compared the levels of lytic toxins and Agr-regulated surface protein, Protein A, between Δ*mroQ* and Δ*agr* mutants (3, 62). We found that toxins leukocidin AB (LukA), Panton-Valentin leukocidin (LukS-PV), and alpha-hemolysin (Hla), were reduced in abundance in a Δ*mroQ* and Δ*agr* mutant, while surface Protein A levels were increased (Fig 4A). Indeed, a prior transposon mutagenesis screen for deficiencies in hemolytic activity, conducted by Fey *et al*, also identified *SAUSA300_1984* as a gene important for Hla production (63). Consistent with these observations, Δ*mroQ* and Δ*agr* mutants exhibited significantly reduced hemolytic activity compared to WT and Δ*mroQ* + *mroQ* strains (Fig 4B) and both strains were similarly attenuated in a murine intradermal infection model (Fig 4C). In summary, MroQ appears to be required for Agr function in some unknown way.

**Figure 4.**
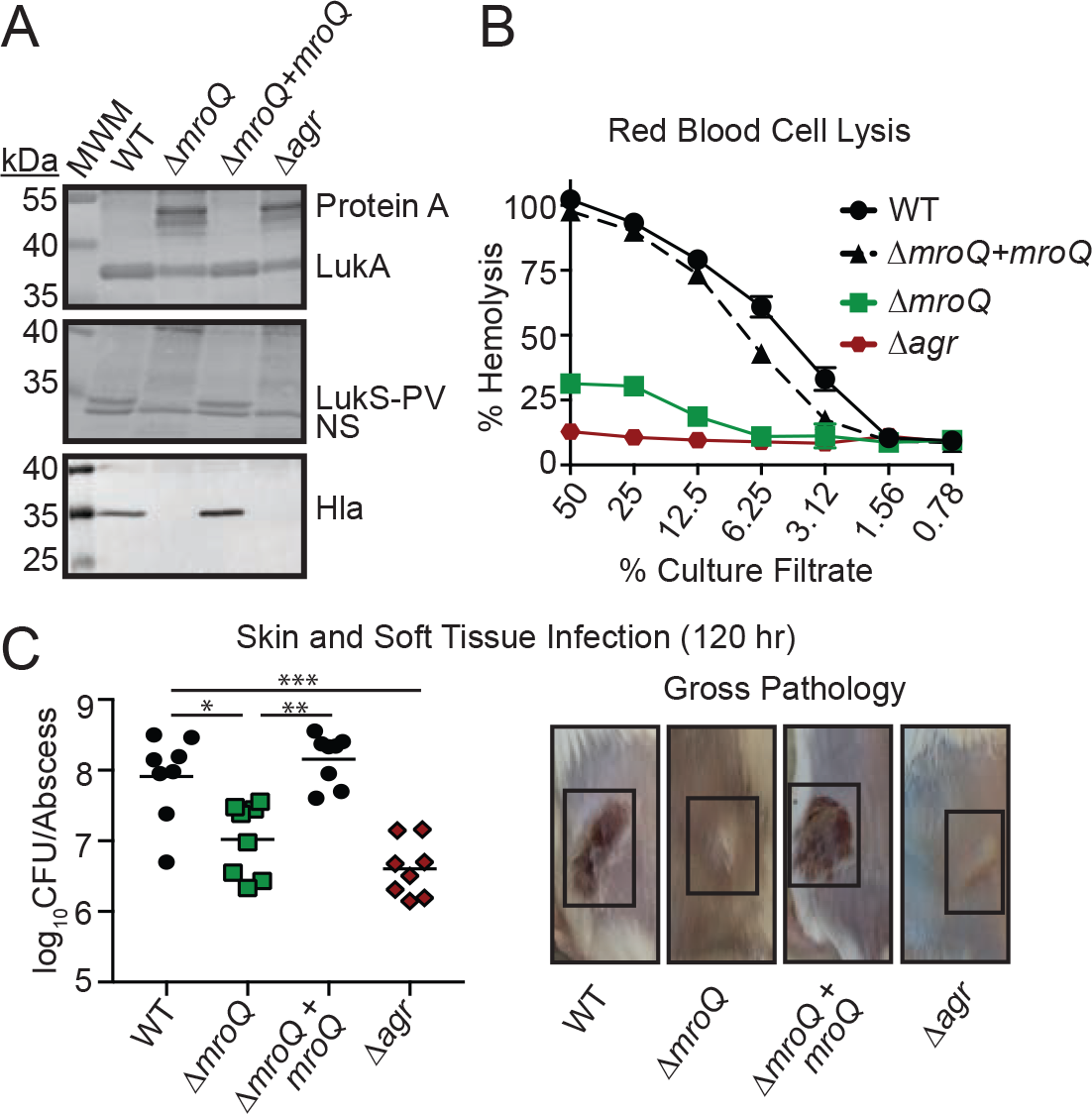
MroQ contributes to Agr function. (A) LukA, LukS-PV, and Hla immunoblots of TCA-precipitated exoproteins from WT, Δ*mroQ*, Δ*mroQ*+*mroQ*, and Δ*agr* strains. Protein A levels were detected based upon binding of α-LukA antibody to Protein A. MWM, Molecular Weight Marker; NS, non-specific band. (B) Rabbit red blood cell lysis of cell free culture filtrates derived from WT, Δ*mroQ*, Δ*mroQ*+*mroQ*, and Δ*agr* strains. (C) Bacterial burden in skin abscesses of mice 120 hours post-infection with WT (n=8), Δ*mroQ* (n=8), Δ*mroQ*+*mroQ* (n=8) and Δ*agr* (n=8) strains. *, *p*<0.05; **, *p*<0.01; ***, *p*<0.001 by non-parametric 1-way ANOVA (Kruskal-Wallis Test) with Dunn’s post-test. Representative images of skin abscesses 120 hours post-infection with WT, Δ*mroQ*, Δ*mroQ*+*mroQ*, and Δ*agr* strains are shown. Hemolysis assay data are from one of at least three experiments conducted in triplicate. Means ± SD are shown (n = 3).

### A Δ*mroQ* mutant is compromised for AIP processing or export

MroQ is a hypothetical type-II CAAX metalloprotease. Despite the clear connections between Agr and *S. aureus* pathobiology, there exists key gaps in our knowledge of the mechanics of Agr system activation (Fig 5A) (5, 64). Most notably: (**i**) the complete range of proteins needed for maturation of AgrD have not been fully elucidated; (**ii**) a dedicated transporter for AIP or its leader peptide has not been conclusively identified; and (**iii**) differential processing of AIPs I-IV has not been investigated. We wondered if the function of MroQ within the Agr system might be to facilitate processing and/or export of the AgrD precursor peptide to generate AIP, or if it promotes function of the histidine kinase AgrC in a manner similar to SpdC and WalK (Fig 5A). To test these possibilities, we first introduced a reporter plasmid that drives the expression of GFP via an AgrA-regulated promoter known as *P3* (pDB59-*P3-gfp*) into Δ*mroQ* and Δ*agr* mutant strains (65). We collected conditioned medium from early stationary phase cultures (AIP production is high) of WT, Δ*mroQ*, Δ*mroQ + mroQ* and Δ*agr* strains and applied them to the reporter strains, followed by measurement of GFP fluorescence. Conditioned medium derived from all strains tested on the Δ*agr* + pDB59-*P3-gfp* reporter strain did not induce promoter activation (Fig 5B). This was expected because the Δ*agr* mutant lacks all components of the Agr system, including those that transmit the AIP signal (AgrC and AgrA). In contrast, when conditioned medium from WT or Δ*mroQ + mroQ* strains was added to the Δ*mroQ* + pDB59-*P3-gfp* reporter strain, we observed a significant increase in fluorescence intensity indicating the AgrC-AgrA signaling axis is intact and able to recognize AIP. However, conditioned medium from Δ*mroQ* and Δ*agr* mutants were unable to induce *P3* promoter activity in this reporter strain background (Fig 5B). Similarly, introduction of the pDB59-*P3-gfp* reporter into Δ*agrB* and Δ*agrB* Δ*mroQ* strains followed by addition of conditioned medium (CM) derived from WT *S. aureus* showed no difference in the ability to transmit signal through AgrC and AgrA (Fig 5C). These data suggest the Agr system defect in a Δ*mroQ* mutant does not occur at the level of AgrC or its ability to propagate signal to AgrA.

**Figure 5.**
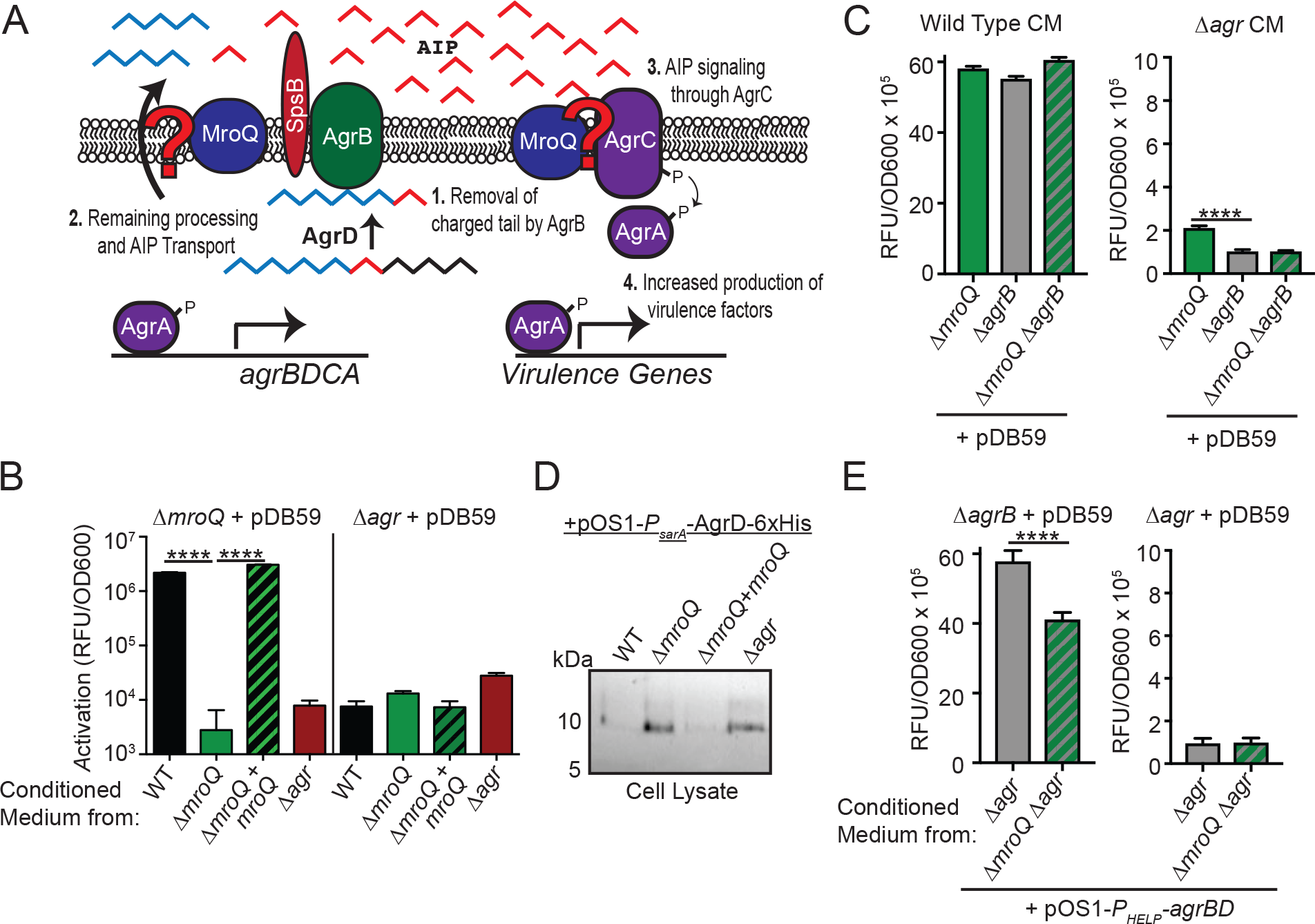
Interrogating effects of MroQ on Agr peptide processing or signaling. (A) Model depicting the potential locations where MroQ might facilitate Agr function. We propose MroQ functions at the level of the peptide-processing module (AgrBD), or at the level of the membrane-embedded histidine kinase (AgrC). (B) *P3-gfp* (pDB59) promoter activation (RFU/OD600) in Δ*mroQ* and Δ*agr* strains upon addition of conditioned medium from WT, Δ*mroQ*, Δ*mroQ*+*mroQ*, and Δ*agr* strains. (C) *P3-gfp* (pDB59) promoter activation (RFU/OD600) in Δ*mroQ*, Δ*agrB*, and Δ*mroQ* Δ*agrB* strains upon addition of conditioned medium (CM) from WT and Δ*agr*. (D) Whole cell lysates of WT, Δ*mroQ*, Δ*mroQ*+*mroQ*, and Δ*agr* strains constitutively expressing AgrD-6x-His (+pOS1-*P*_*sarA*_-*agrD-6x-His*) followed by immunoblotting with α-His monoclonal antibody to detect full-length (unprocessed) AgrD. (E) *P3-gfp* (pDB59) promoter activation (RFU/OD600) in Δ*agrB*, and Δ*agr* strains upon addition of conditioned medium from Δ*agr* and Δ*agr* Δ*mroQ* constitutively expressing *agrBD* (+ pOS1-*P*_*HELP*_-*agrBD*). ****, *p*<0.0001 by 1-way ANOVA with Tukey’s post-test (C) and two-tailed T-test (E). Data shown are from one of at least three experiments conducted in triplicate. Means ± SD are shown (n = 3).

The absence of a clear effect of MroQ on AgrC signaling lead us to interrogate whether or not MroQ affects AgrD protein processing/export. We generated a plasmid that expresses AgrD-6xHis under the control of a constitutive promoter (pOS1-*P*_*sarA*_-*agrD-6xHis*) (66, 67). The plasmid was introduced into WT, Δ*mroQ*, Δ*mroQ+mroQ*, and Δ*agr* strains. It is known that unprocessed AgrD accumulates in the absence of the AgrB peptidase, indicating that processing of AgrD by AgrB precedes export (3, 27–30). We hypothesized that if MroQ is involved in promoting an AgrD processing event similar to that of AgrB, then we should detect full length AgrD-6xHis in the Δ*mroQ* mutant. Indeed, full length AgrD-6xHis was detectable when expressed in the Δ*mroQ* and Δ*agr* strains (Fig 5D). To further determine if MroQ is involved in regulating AgrBD functions related to AIP processing/export, we generated a plasmid that constitutively expresses *agrBD* driven by the synthetic promoter, *P*_*HELP*_, and introduced this plasmid (pOS1-*P*_*HELP*_-*agrBD*) into Δ*agr* or Δ*agr* Δ*mroQ* strains followed by determination of whether or not conditioned medium derived from these strains is able to activate the pDB59-*P3-gfp* reporter in a Δ*agrB* or Δ*agr* mutant background. We found that conditioned medium derived from the Δ*agr* Δ*mroQ* + pOS1-*P*_*HELP*_-*agrBD* exhibited a 30% reduction in P3 promoter activity compared to conditioned medium derived from the Δ*agr* + pOS1-*P*_*HELP*_-*agrBD* strain (Fig 5E). These data support the hypothesis that MroQ plays a role in promoting the functionality of the AgrB-AgrD peptide processing/export axis of the Agr system.

### Conserved active site amino acids are required for MroQ regulation of the Agr system

The type-II CAAX protease family of enzymes is conserved in both prokaryotes and eukaryotes. In eukaryotes, proteolytic activity occurs at a CAAX motif on the C-terminal end of target proteins, ultimately leading to lipidation. In bacteria, the functions of the CAAX protease family members are not as clear, but some proteins in this family use proteolytic activity to confer immunity against bacteriocins (68), while others engage in intramembrane proteolysis events to promote adaptive traits (52, 69). Of the three additional characterized CAAX protease enzymes in *S. aureus* (SpdA-C), the proteolytic activity of the enzyme does not appear to be required for function. This is also true of the Abx1 enzyme of Group B *Streptococcus*. Thus, the CAAX proteases of bacteria appear to have diverse functions, some of which depend on proteolytic activity and others that do not. To test if the active site of MroQ is required to promote AgrBD activity, we introduced mutations into the conserved CAAX protease amino acid residues within the EEXXXR and FXXXH motifs previously determined to be required for enzymatic activity in a range of type-II CAAX proteases (50, 52, 68, 70, 71). Alanine substitutions were introduced at E141, E142, and H180 followed by expression of point mutants in single copy under the control of the *mroQ* native promoter in Δ*mroQ S. aureus* (Fig. 6A). As evidenced by immunoblot analysis, the expression of MroQ(E141A) was unable to complement alpha toxin secretion defects, whereas MroQ(E142A) and MroQ(H180A) exhibited some restoration in alpha toxin production by immunoblot, with MroQ(E142A) supernatant containing greater amounts of alpha toxin than MroQ(H180A) (Fig. 6B). The observed differences in alpha toxin levels by immunoblot were recapitulated in hemolytic activity assays where MroQ(E142A) exhibited partial restoration in hemolytic activity on rabbit red blood cells, but MroQ(E141A) and MroQ(H180A) did not (Fig. 6C). Consistent with the observed effects on alpha toxin secretion and hemolytic activity, MroQ(E142A) partially complemented protein A production defects (Fig. 6D). In contrast, none of the three mutations restored LukS or LukA production to an observable degree by immunoblot and the MroQ point mutants failed to complement changes in macrophage activation caused by supernatant derived from a Δ*mroQ* mutant (Fig. 6D-E). Together, these data suggest that conserved amino acids implicated in type-II CAAX protease activity are important for the effects of MroQ on AgrBD function. Residue E141 is required for MroQ function, whereas residues E142 and H180 promote optimal activity.

**Figure 6.**
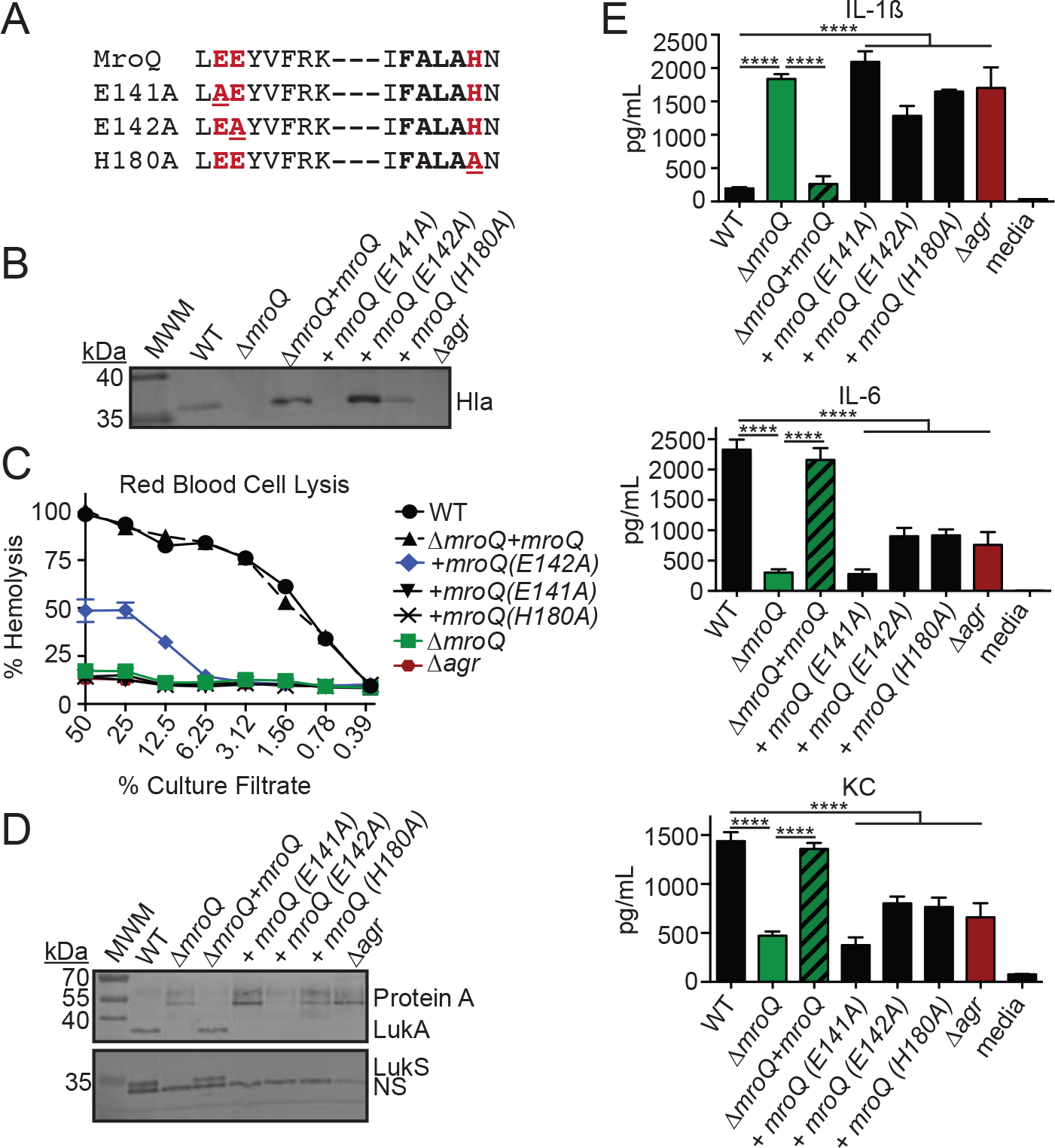
MroQ active site residues are required for regulation of Agr by MroQ. (A) Amino acid sequences of the EEXXXR and FXXXH motifs that comprise the Type-II CAAX protease active site. Locations of site-directed amino acid substitutions are underlined. (B) Hla immunoblots of TCA-precipitated exoproteins from WT, Δ*mroQ*, Δ*mroQ*+*mroQ*, Δ*mroQ*+*mroQ(E141A)*, Δ*mroQ*+*mroQ(E142A)*, Δ*mroQ*+*mroQ(H180A)*, and Δ*agr* strains. (C) Rabbit red blood cell lysis of cell free culture filtrates derived from WT, Δ*mroQ*, Δ*mroQ*+*mroQ*, Δ*mroQ*+*mroQ(E141A)*, Δ*mroQ*+*mroQ(E142A)*, Δ*mroQ*+*mroQ(H180A)*, and Δ*agr* strains. (D) LukA and LukS-PV immunoblots of TCA-precipitated exoproteins from WT, Δ*mroQ*, Δ*mroQ*+*mroQ*, Δ*mroQ*+*mroQ(E141A)*, Δ*mroQ*+*mroQ(E142A)*, Δ*mroQ*+*mroQ(H180A)*, and Δ*agr* strains. Protein A levels were detected based upon binding of α-LukA antibody to Protein A. MWM, Molecular Weight Marker; NS, non-specific band. (E) IL-1β, IL-6, and KC production (pg/mL) by BMM after addition of supernatant from WT, Δ*mroQ*, Δ*mroQ*+*mroQ*, Δ*mroQ*+*mroQ(E141A)*, Δ*mroQ*+*mroQ(E142A)*, Δ*mroQ*+*mroQ(H180A)*, and Δ*agr* strains grown in TSB. ****, *p*<0.0001 by 1-way ANOVA with Tukey’s post-test. Data shown are from one of at least three experiments conducted in triplicate. Means ± SD are shown (n = 3).

### Investigation of the role of MroQ amino acid residues E141, E142, and H180 in Agr-mediated gene expression and virulence

We next sought to determine the extent with which Δ*mroQ* mutant strains expressing MroQ(E141A), MroQ(E142A), and MroQ(H180A) alter gene expression and pathogenesis. We first conducted P3 promoter activation analyses to evaluate whether or not generation and release of functional AIP occurs in strains expressing each point mutant. As anticipated, conditioned medium derived from Δ*mroQ* + MroQ(E141A) and Δ*mroQ* + MroQ(H180A) were unable to appreciably activate the P3-gfp promoter fusion above background levels when applied to the Δ*mroQ* + pDB59 reporter strain (Fig 7A). In contrast, conditioned medium derived from Δ*mroQ* + MroQ(E142A) led to partial activation with a greater than 10-fold increase in fluorescence intensity compared to background levels (Fig. 7A). When Δ*mroQ* + MroQ(E141A), Δ*mroQ* + MroQ(E142A), Δ*mroQ* + MroQ(H180A) were used in an intradermal infection model, we found that Δ*mroQ* + MroQ(E141A) exhibited a 10-fold reduction in CFU, similar to a Δ*mroQ* and Δ*agr* mutant. In contrast, the Δ*mroQ* + MroQ(E142A), and Δ*mroQ* + MroQ(H180A) strains restored infection to near WT levels (Fig. 7B). Thus, MroQ(E141A) appears to be the most crucial active site mutation as it is necessary for MroQ function in vitro and in vivo, whereas MroQ(E142A) and MroQ(H180A) confer near normal virulence characteristics during skin and soft tissue infection, an outcome consistent with their partial restoration of in vitro defects.

**Figure 7.**
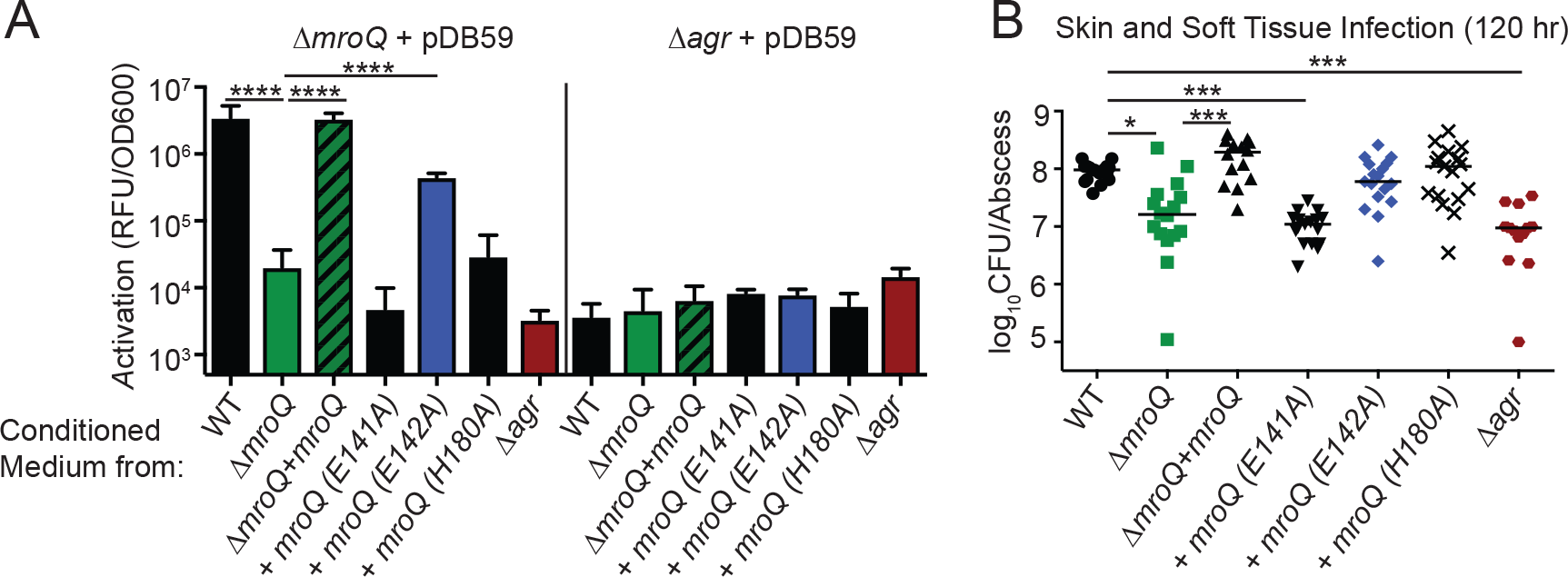
MroQ amino acid substitution E141A is required for Agr activation and full virulence. (A) *P3-gfp* (pDB59) promoter activation (RFU/OD600) in Δ*mroQ* and Δ*agr* strains upon addition of conditioned medium from WT, Δ*mroQ*, Δ*mroQ*+*mroQ*, Δ*mroQ*+*mroQ(E141A)*, Δ*mroQ*+*mroQ(E142A)*, Δ*mroQ*+*mroQ(H180A)*, and Δ*agr* strains. Data shown are from one of at least three experiments conducted in triplicate. Means ± SD are shown (n = 3). ****, *p*<0.0001 by 1-way ANOVA with Tukey’s post-test. (B) Bacterial burden in skin abscesses of mice 5 days post-infection with WT (n=16), Δ*mroQ* (n=16), Δ*mroQ*+*mroQ* (n=14), Δ*mroQ*+*mroQ(E141A)* (n=17) (Δ*mroQ*+*mroQ(E142A)* (n=17), Δ*mroQ*+*mroQ(H180A)* (n=17), and Δ*agr* (n=16). *, *p*<0.05; ****, *p*<0.001 by non-parametric 1-way ANOVA (Kruskal-Wallis Test) with Dunn’s post-test.

## Discussion

Many Gram-positive pathogens use small peptide signaling pathways to control the expression of virulence genes. In this work, we present the discovery of a membrane-embedded protease that plays a major role in promoting the function of the Agr system of *S. aureus*. Our data suggest that the abortive infectivity protein (Abi), or type-II CAAX peptidase, SAUSA300_1984 (MroQ), directly interfaces with the Agr system to promote virulence factor gene expression and pathogenesis in a murine skin and soft tissue infection model. Unlike other type-II CAAX proteases of *S. aureus* and Group B *Streptococcus* (55, 57, 58), MroQ requires its conserved active site amino acids to confer normal functionality to the Agr system. Our studies to probe the location in the Agr pathway where MroQ acts indicate that the AgrC/A signaling axis remains functional and that the AgrBD axis, involved in peptide maturation, is likely compromised. Together, these findings expand the view of peptide-based signaling in *S. aureus* and highlight a newly uncovered regulatory input into virulence factor gene expression that dramatically affects virulence outcomes. Gram-positive pathogens of urgent clinical importance including *Clostridioides difficile* and *Enterococcus faecalis*, as well as significant threats to food safety such as *Listeria monocytogenes* each harbors Agr systems that contribute to disease (72–78). Thus, this work provides insights into alternative pathways of regulation of the Agr system that may extend to other Gram-positive microbes of clinical importance.

Our studies have determined that the secreted factors derived from either a Δ*mroQ* or Δ*agr* mutant cause significant release of IL-1β, but low levels of other inflammatory cytokines and chemokines from primary murine macrophages in culture (Fig 1). The production of the inflammatory cytokine IL-1β in superficial tissues is known to be critical for controlling skin and soft tissue infection caused by *S. aureus* (79–83). In contrast, the production of IL-1β during systemic infection is associated with increased immunopathology (84, 85). Agr-defective isolates of *S. aureus* are routinely isolated from patients with bacteremia (64). The reason these Agr-defectives arise and their association with morbidity in bacteremia is not well understood; however, the observation is recapitulated in murine infection models wherein *agr* mutants persist in deep tissues during bacteremia (86). Some potential mechanisms behind these observations have been proposed, including a role for *agr*-defectives in development of a persister state, as well as disruption of the optimal immune environment by *agr*-defective strains (64). Although a Δ*mroQ* mutant does not elicit increased production of IL-1β compared to WT at the time points tested in vivo, we postulate that the reduced levels of pro-inflammatory cytokines and maintenance of IL-1β may be linked to improved disease outcomes (Fig 2). The possibility that the inflammatory shifts imparted by a loss of Agr functionality might dictate clearance or survival in superficial versus systemic infection is currently being interrogated by our laboratory.

The mechanics of cyclic AIP maturation by *S. aureus* is not fully understood (5). Generation of AIP absolutely requires C-terminal cleavage and formation of a thiolactone ring by the Agr locus enzyme, AgrB (27–30). The events that occur after thiolactone ring formation remain ambiguous, but at minimum must involve additional proteins for proteolytic cleavage and transport (5, 32). Our data demonstrates that the putative metalloprotease, MroQ, does not seem to influence the ability of the histidine kinase AgrC to transmit signal to AgrA (Fig 5C). Thus, it does not appear that MroQ is directly regulating the activity of AgrC. This observation is different from that observed for Abx1 from Group B *Streptococcus* or SpdC of *S. aureus*, which are implicated in the regulation of CovS and WalK histidine kinases respectively (57, 58). These studies used bacterial two-hybrid assays to determine that Abx1 and SpdC interact with WalK and CovS and that deletion or over-expression changes kinase activity. Aside from these physical associations, no additional evidence was provided to describe how each Abi protein was mediating its function, but the studies do note that SpdC negatively regulates WalK and overexpression of Abx1 promotes CovS activity (57, 58). Uncoupling *agrBD* gene expression from its auto-regulatory circuit has allowed us to identify a defect in AIP peptide production when MroQ is absent (Fig 5E). We find that conditioned medium derived from Δ*agr* + pOS1-*P*_*HELP*_-*agrBD* is better able to activate a *P3-gfp* reporter compared to conditioned medium derived from Δ*agr* Δ*mroQ* + pOS1-*P*_*HELP*_-*agrBD*. Importantly, the absence of MroQ does not eliminate reporter activation, suggesting that either overexpression of *agrBD* can compensate, or that MroQ activity on AgrBD is not essential for peptide maturation/release, rather it improves its efficiency in some way. Thus, MroQ appears to behave differently from Abx1 and SpdC, where it acts to promote functionality of the peptide maturation module of Agr and is important for the optimal generation or export of AIP in the Type I Agr MRSA strain LAC. We do not yet know precisely how MroQ is mediating its effects on the AgrBD module, however a Δ*mroQ* mutant does not release appreciable amounts of active AIP (Fig 5 and 7) and harbors increased amounts of unprocessed cytosolic AIP (Fig 5D). It is notable that although a Δ*mroQ* mutant closely resembles a Δ*agr* mutant, there appears to be some low-level intrinsic activation of the Agr system in a Δ*mroQ* mutant as evidenced by P3 promoter activity that rises above levels of a Δ*agr* mutant (Figs 5C and 7A). This observation indicates that mature AIP is at least capable of being produced by a Δ*mroQ* mutant, though at low levels. Taken together, we propose MroQ promotes efficient AgrD proteolytic processing, transport, or both. This may occur through direct effects on AgrB function or on the AgrD peptide itself. We are currently exploring these possibilities.

*S. aureus* strains encode one of four allelic variants of AIP, all of which are capable of causing clinical disease in humans (36–40). Our data indicate that MroQ is critical for Agr activity of a Type I strain. AgrB, the protease that cleaves the C-terminus of AgrD, has considerable sequence variability among Agr types, rendering C-terminal cleavage specific to the four Agr groups (3, 30, 33). There is negligible sequence variation at the amino acid level for MroQ in strains that harbor each of the four Agr alleles (data not shown). Therefore, we suspect that either the activity of MroQ on the Agr system is broad and not influenced by Agr Type, or that alternative enzymes, perhaps other CAAX proteases, modulate the functions of different Agr types. Given the strict dependency on the length and amino acid composition of the N-terminal tails of AIP for signaling and given that precise peptide cleavage at the N-terminus is a requisite for activity, we favor the possibility that additional peptidases may promote full maturation or export of these AIP sequence variants (39).

One important difference noted in the function of MroQ, compared to that of other type-II CAAX proteases in *S. aureus* is that conserved active site residues appear to be important for the activity of MroQ, but not for the CAAX proteases SpdA-C, or Group B *Streptococcus* Abx1. This represents an additional departure from the models put forth by these CAAX proteases and suggests that MroQ may alternatively function in a way that is similar to that of proteases whose activities involve peptide hydrolysis or intramembrane proteolysis (50, 52, 68, 70, 71). Thus far we know that the glutamate at position 141 is critical for MroQ function, whereas mutations of glutamate at position 142 and the histidine at position 180 do not confer complete null phenotypes. In other Abi proteins, the second glutamic acid in the EEXXXR motif is usually essential for activity, an observation that is contrary to what was observed for MroQ. While E142A and H180A mutations confer a partial restoration in Agr activity, only the E141A point mutation is attenuated in vivo to the same level as a Δ*mroQ* or Δ*agr* mutant during skin and soft tissue infection. This outcome suggests that partial Agr activity conferred by expression of MroQ(E142A) and MroQ(H180A), but not MroQ(E141A), is sufficient to confer normal infection kinetics.

Altogether, this work has expanded our understanding of the regulatory inputs that dictate cyclic peptide quorum sensing in *S. aureus*. Our studies represent an expansion of the Agr regulatory circuit that may have broad impacts on therapeutic design for non-complicated infection (12, 14, 87–89). Further, the work highlights the complexities of peptide based signaling and the adaptive traits that promote virulence in *S. aureus*. Our knowledge of the activities of proteins from the Abi protein family in Gram-positive pathogens remains in its infancy. This work adds to the growing compendium of work on this protein class and their diverse range of functions, some of which are crucial for Gram-positive bacterial virulence.

## Materials and Methods

### Bacterial strains and culture conditions

*S. aureus* LAC (AH-1263) was used as the wild type (WT) strain for the experiments in this study. LAC (AH-1263) is a *S. aureus* USA300 clinical isolate cured of its plasmids (90). All other bacterial strains used in this work are described in Table 1. Most recombinant plasmids were passaged through *E. coli* DH5α before propagation in *S. aureus*. When constructing complementation plasmids harboring the *mroQ* gene, we found that *mroQ* exhibited signs of toxicity in *E. coli*, resulting in a high frequency of secondary mutations in the coding sequence. As an alternative, *E. coli* BH10C was used as a host strain; BH10C reduces copy number of plasmids to near single-copy levels to limit the abundance of potentially toxic genes in *E. coli* and reduces the likelihood of mutations (91). After passage through *E. coli, S. aureus* RN4220 and RN9011 (RN4220+pRN7023) (61) were used as intermediate strains, followed by electroporation or transduction into AH-1263 or its isogenic mutant derivatives. All *E. coli* strains were grown in Lysogeny Broth (LB) (Amresco), while *S. aureus* strains were grown in either Tryptic Soy Broth (TSB) (Criterion) or Roswell Park Memorial Institute medium (RPMI) (Corning) supplemented with 1% casamino acids (Amresco) and 2.4 mM Sodium bicarbonate (Amresco). When necessary, media were supplemented with antibiotics at the following concentrations: ampicillin (Amp), 100 μg/mL; erythromycin (Erm), 5 μg/mL; chloramphenicol (Cm), 10 μg/mL; anhydrous tetracycline (AnTet) (Acros Organics), 1 μg/mL; and tetracycline (Tet) at 10 μg/mL (Amresco and Acros Organics). Cadmium chloride was used at 0.1-0.3 mM to select for pJC1111 transductants. Bacterial growth was monitored using a Genesys 10S UV-Vis spectrophotometer by measuring culture optical density at 600 nm (OD600) (Thermo).

**Table 1.**
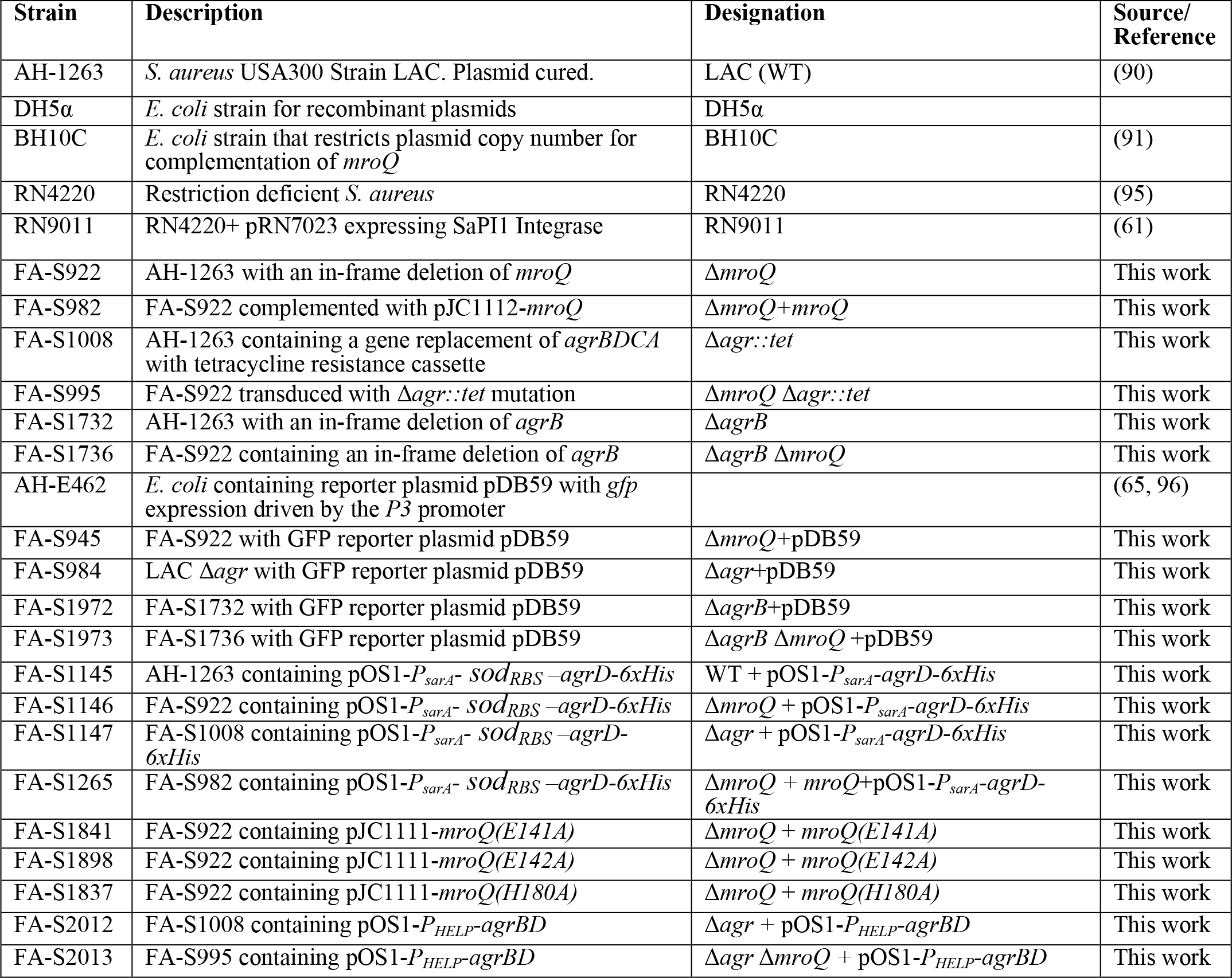
List of strains used in this study.

### Molecular genetic techniques

To isolate genomic DNA from *S. aureus*, cultures were grown overnight in 5 mL TSB at 37°C with shaking at 200 rpm. 1.5 mL of bacteria was pelleted by centrifugation and resuspended in 200 μL of TSM buffer (50 mM Tris, 0.5 M Sucrose, 10 mM MgCl2 (pH 7.5)) and 2.5 μL of a lysostaphin (Ambi Products) stock (2 mg/mL) to achieve a final concentration of 25 μg/mL. Samples were incubated at 37°C for 15 minutes to digest the cell wall of *S. aureus*, followed by centrifugation at 14000 rpm for 5 minutes to pellet bacteria. Supernatants were discarded and the genomic DNA was isolated using a Wizard Genomic DNA purification kit (Promega). PCR was performed in a FlexID Mastercycler (Eppendorf) using Phusion High-Fidelity DNA Polymerase (New England Biolabs) or GoTaq DNA Polymerase (Promega) and dNTPs (Quanta BioSciences). For all PCR reactions oligonucleotides were ordered from Eurofins (see Table 2). Electrophoresis of DNA samples was performed in 0.8% agarose (Amresco) gels. DNA digestions were performed using the following restriction endonucleases: XhoI, BamHI, EcoRI, KpnI, XbaI, SacI and PstI (New England Biolabs) following the manufacturer’s suggested protocols and all digested plasmids were subsequently treated with Shrimp Alkaline Phosphatase (Amresco). Ligations were performed using T4 DNA Ligase (New England Biolabs) and were incubated in an Eppendorf ThermoMixer at 16°C overnight. When necessary, PCR purification and DNA gel extraction was carried out using QIAGEN QIAquick kits.

**Table 2.**
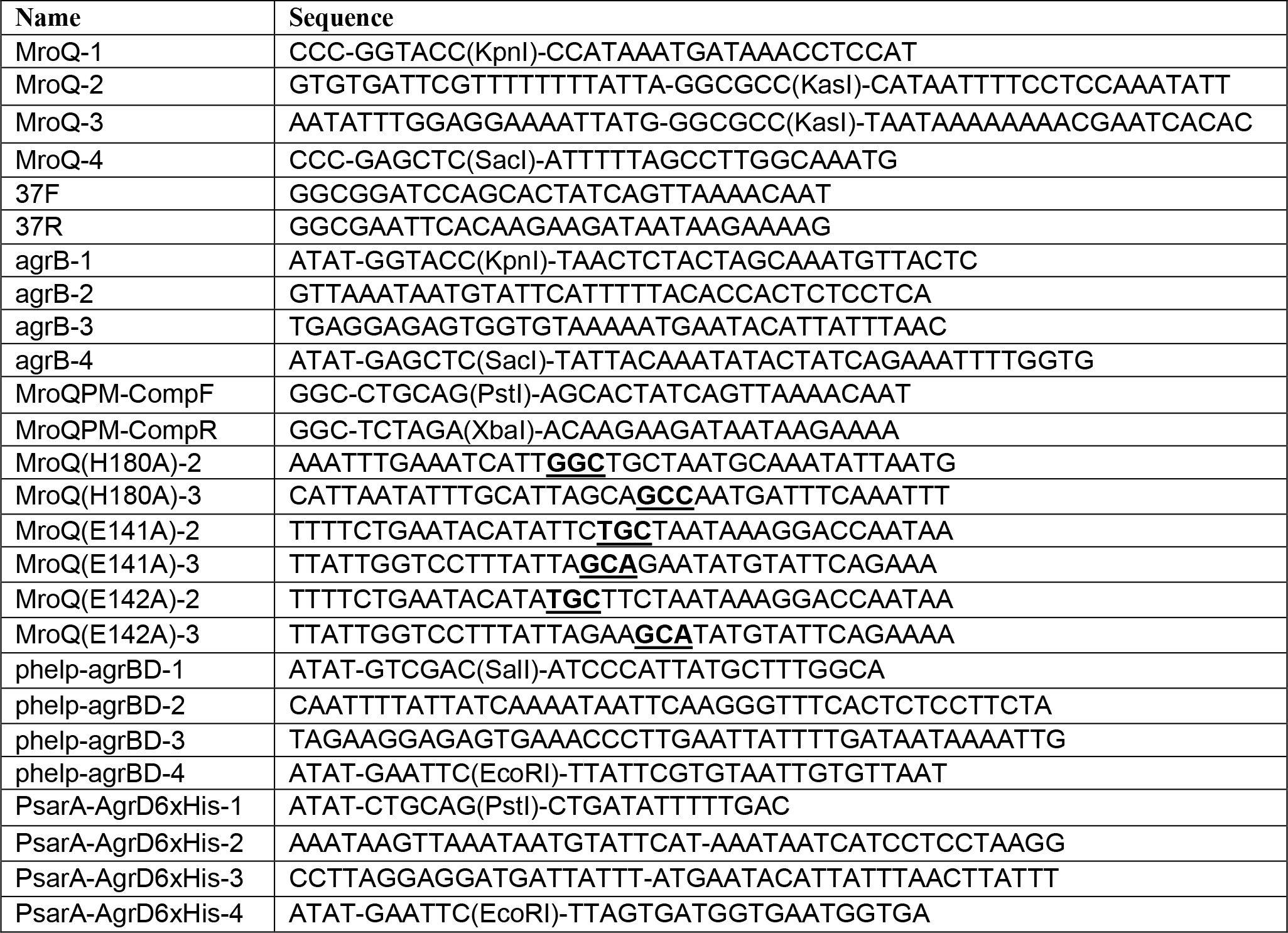
List of oligonucleotides used in this study.

To isolate plasmids from *S. aureus*, 5 mL overnight cultures of bacteria were grown in TSB at 37°C with shaking at 200 rpm. The next day, bacteria were pelleted by centrifugation at 4200 rpm for 15 minutes and resuspended in 400 μl TSM buffer containing 20 μl of lysostaphin stock solution (2 mg/mL) to achieve a final concentration of 0.1 mg/mL. Samples were incubated for 10 minutes at 37°C to digest the cell wall, followed by centrifugation at 13000 rpm for 2 minutes to pellet the bacterial cells. Thereafter, a Qiagen miniprep kit was used to isolate plasmid DNA. DNA was eluted with sterile water in all cases. DNA sequencing was performed by Genscript or Genewiz.

### Bacteriophage-mediated generalized transduction

Transduction was used to transfer stably integrated complementation plasmids between strains, as well as to mobilize marked mutations within the *S. aureus* chromosome (Δ*agr*::*tet*). *S. aureus-*specific bacteriophages ϕ11 or 80α were used in this study. Donor strains were grown overnight in 3 mL of TSB/LB (1:1) supplemented with 5 mM calcium chloride (CaCl_2_) (Amresco) and 5 mM magnesium sulfate (MgSO_4_) (Amresco) at 37°C with shaking at 200 rpm. The next day, overnight cultures were diluted 1:100 in 10 mL fresh TSB/LB (1:1) supplemented with 5 mM CaCl_2_ and 5 mM MgSO_4_. Samples were grown for ~3 hours at 37°C with shaking at 200 rpm until reaching an OD600 of 0.3 - 0.9. To package donor DNA into transducing phage, 100 μL of 10-fold serial dilutions (10^−1^ to 10^−10^) of bacteriophage stock in TMG (10 mM Tris pH 7.5, 5 mM MgCl_2_, 0.01% gelatin (v/v)) were added to 15 mL conical tubes and incubated with 500 μL of the donor *S. aureus* culture. The phage-bacteria mixture was gently vortexed and incubated at room temperature for 30 minutes. 2.5 mL of melted and cooled CY top agar (casamino acids 5 g/L, yeast extract 5 g/L, glucose 5 g/L, NaCl 6 g/L, 7.5 g/L agar) supplemented with 5 mM CaCl_2_ and 5 mM MgSO_4_ was added to the bacteria-phage mixture and immediately poured onto TSA plates followed by incubation overnight at 30°C. The following day, phage were harvested from two to three plates with confluent plaques. Bacteriophage stocks were stored at 4°C.

To transduce plasmids and marked mutations, recipient strains were first grown overnight in 20 mL of TSB/LB (1:1) supplemented with 5 mM CaCl_2_ at 37°C. Samples were then centrifuged for 15 minutes at maximum speed, supernatants were discarded, and bacterial pellets suspended in 3 mL of CY medium (casamino acids 5 g/L, yeast extract 5 g/L glucose 5 g/L NaCl 6 g/L) supplemented with 5 mM CaCl_2_. Four infection conditions were set up with either bacteria alone (mock) or 10-fold serial dilutions of bacteria infected with 100 μL of bacteriophage stock containing donor DNA. Tubes were mixed by inverting every ten minutes for 30 minutes at room temperature, followed by the addition 40 mM sodium citrate to each tube and incubation for 30 minutes at room temperature. Tubes were centrifuged at maximum speed for 5 minutes and washed twice in 500 μL of CY medium supplemented with 40 mM sodium citrate, followed by resuspension in 200 μL of CY medium supplemented with 40 mM sodium citrate and spreading onto CY agar or TSA plates containing 10 mM sodium citrate and any antibiotics necessary for selection. Plates were placed at 37°C overnight. Potential transductants were screened for antibiotic resistance and acquisition of desired mutations by PCR and DNA sequencing.

### Generation of *mroQ* and *agrB* in-frame deletion mutants

*mroQ* and *agrB* in-frame deletion mutants were generated using the temperature-sensitive mutagenesis plasmid pIMAY (92). For *mroQ*, two fragments that correspond to ~500 base pair regions of sequence homology immediately upstream and downstream of *mroQ* were amplified. Oligonucleotides MroQ-1 and MroQ-2 were used to amplify the region upstream of *mroQ*, while oligonucleotides MroQ-3 and MroQ-4 were used to amplify the region downstream of *mroQ* (Table 2). The fragments were joined by splicing by overlap extension (SOEing) PCR using oligonucleotides MroQ-1 and MroQ-4 and subcloned into the multi-cloning site of pIMAY using restriction endonucleases KpnI and SacI. For *agrB*, two fragments that correspond to ~500 base pair regions of sequence homology immediately upstream and downstream of *agrB* were amplified. Oligonucleotides agrB-1 and agrB-2 were used to amplify the region upstream of *agrB*, while oligonucleotides agrB-3 and agrB-4 were used to amplify the region downstream of *agrB* (Table 2). The fragments were joined by splicing by overlap extension (SOEing) PCR using oligonucleotides agrB-1 and agrB-4 and subcloned into the multi-cloning site of pIMAY using restriction endonucleases KpnI and SacI. Mutagenesis was carried out according to our previously published protocols (48, 49). PCR and DNA sequencing analysis were used to determine successful generation of mutants.

### Generation of Δ*mroQ* +*mroQ*, Δ*mroQ* +*mroQ(E141A)*, Δ*mroQ* +*mroQ(E142A)*, and Δ*mroQ* +*mroQ(H180A)* complementation strains

A single copy chromosomal complementation strain driving *mroQ* expression under its native promoter was constructed using the integrative plasmid pJC1112 (61). The pJC1112-*mroQ* plasmid was generated by PCR using oligonucleotides 37F and 37R (Table 2). The resulting amplicon was then subcloned into the pJC1112 vector using EcoRI and BamHI restriction endonucleases followed by transformation into BH10C *E. coli* and electroporation into the RN9011 where the plasmid integrates at the SapI1 site. The integrated complementation vector was then transduced into a Δ*mroQ* mutant as described above. Successful introduction of the complementation construct was determined by PCR and sequencing. To generate complementation strains harboring amino acid substitutions in MroQ catalytic residues, primer MroQPMComp-F was paired with MroQ(E141A)-2, MroQ(E142A)-2, or MroQ(H180A)-2 and primer MroQPMComp-R was paired with MroQ(E141A)-3, MroQ(E142A)-3, or MroQ(H180A)-3. The resulting amplicons from these two PCRs were used in a SOEing PCR with primers MroQ-PMComp-F and MroQ-PMComp-R followed by subcloning the product into pJC1111 using XbaI and PstI restriction endonucleases followed by transformation into BH10C *E. coli* and electroporation into the RN9011 where the plasmid integrates at the SapI1 site. The integrated complementation vector was then transduced into a Δ*mroQ* mutant as described above.

### Construction of Δ*agr*::*tet* mutants

A marked deletion mutation of *agrBDCA* (Δ*agr*::*tet*) was transduced into AH-1263 (WT), Δ*mroQ*, and Δ*agrB* by bacteriophage-mediated transduction as described above.

### Construction of *P3-gfp* reporter strains

The pDB59 reporter plasmid contains the *P3* promoter of the Agr system driving the expression of *gfp* (56). The plasmid was isolated from *E. coli* and passaged through RN4220, followed by electroporation into WT, Δ*mroQ*, Δ*agrB*, Δ*agrB* Δ*mroQ*, and Δ*agr::tet*.

### Construction of pOS1-*P*_*sarA*_-*agrD-6xHis* expression plasmid

An AgrD-6x-His expression plasmid was generated by fusing the *P*_*sarA*_ promoter linked to the *S. aureus* superoxide dismutase (*sod)* ribosomal binding site (66, 67) and to the coding sequence of AgrD harboring a 6x-His tag on its C-terminus. *P*_*sarA*_-*sod*_*RBS*_ was amplified using primers PsarA-AgrD6xHis-1 and PsarA-AgrD6xHis-2 and agrD-6xHis was amplified using oligonucleotides PsarA-AgrD6xHis-3 and PsarA-AgrD6xHis-4. A SOEing PCR was performed to splice these amplicons together using primers PsarA-AgrD6xHis-1 and PsarA-AgrD6xHis-4. The resulting amplicon was cloned into pOS1 using PstI and EcoRI restriction endonucleases and the product was transformed into DH5α, passaged through RN4220, and electroporated into wild type, Δ*mroQ*, Δ*mroQ+mroQ*, and Δ*agr* isogenic strains.

### Construction of pOS1-*P*_*HELP*_-*agrBD* plasmid

An *agrBD* expression plasmid was generated by fusing the *P*_*HELP*_ promoter (48, 49, 93) to *agrBD*. *P*_*HELP*_ was amplified using primers phelp-AgrBD-1 and phelp-AgrBD-2 and *agrBD* was amplified using oligonucleotides phelp-AgrBD-3 and phelp-AgrBD-4. A SOEing PCR was performed to splice the resulting amplicons together using primers phelp-AgrBD-1 and phelp-AgrBD-4. The final amplicon was cloned into pOS1 using SalI and EcoRI restriction endonucleases and the resulting plasmid was transformed into BH10C *E. coli*, passaged through RN4220, and electroporated into wild type, Δ*agr*, and Δ*agr* Δ*mroQ* isogenic strains.

### Whole cell lysate preparation

5 mL of *S. aureus* strains were grown overnight at 37°C with shaking at 200 rpm. Overnight cultures were diluted 1:100 in 20 mL of fresh medium and allowed to replicate for an additional 8 hours at 37°C with shaking at 200 rpm. After 8 hours, the OD600 was measured and samples were centrifuged at 4200 rpm for 15 minutes to pellet the bacteria. Supernatants were removed and the bacterial cell pellets were stored at −80°C until use. Pellets were thawed on ice and resuspended in 250 μL of PBS supplemented with 1 M NaCl and 6 M Urea (VWR). The resuspended bacteria were then added to screw cap microcentrifuge lysing tubes (Fisher Scientific) containing 250 μL 0.1 mm glass cell disruption beads (Scientific Industries, Inc.). Cells were lysed using a Fast Prep-24 5G (MP Biomedicals) at speed 5.0 for 20 seconds, subsequent incubation on ice for 5 minutes, and then disrupted a second time at speed 4.5 for 20 seconds. 50 μL of 6x SDS Sample buffer (0.375 M Tris pH 6.8, 12% SDS, 60% glycerol, 0.6 M DTT, 0.06% bromophenol blue) was added directly to lysed cells and boiled for 10 minutes, followed by centrifugation at maximum speed for 15 minutes at 4°C. Samples were stored at −20°C for no longer then a few days.

### Exoprotein preparations

*S. aureus* strains were grown at 37°C with shaking at 200 rpm overnight. Overnight cultures were diluted 1:100 in fresh medium and allowed to grow to stationary phase at 37°C with shaking at 200 rpm (~8 hours). OD600 was measured and samples were centrifuged at 4200 rpm for 15 minutes. 1.3 mL of supernatants were collected in 1.5 mL microcentrifuge tubes and cooled on ice for approximately 30 minutes followed by addition of 150 μl of 100% trichloroacetic acid (TCA) and overnight incubation at 4°C. The next 24 day, samples were centrifuged at maximum speed for 15 minutes at 4°C and supernatants were removed and discarded. Precipitated proteins were washed with 1 mL of 100% ethanol (Decon Laboratories Inc.) and incubated at 4°C for 30 minutes, followed by a 15-minute spin at maximum speed at 4°C. Protein pellets were air dried at room temperature for about one hour, followed by dissolving the pellet in TCA-SDS sample buffer (0.5 M Tris-HCl buffer, 4%SDS mixed 1:1 with 2x SDS Sample buffer) and boiling for 10 minutes before storage at −20°C.

### Immunoblot for secreted virulence factors

Wild type, Δ*mroQ*, Δ*mroQ+mroQ*, Δ*mroQ+mroQ(E141A)*, Δ*mroQ+mroQ(E142A)*, Δ*mroQ+mroQ(H180A)*, and Δ*agr* strains were grown in 5 mL of TSB for eight hours at 37°C with shaking at 200 rpm and supernatants were collected for TCA precipitation as described above. Prior to loading, protein samples were normalized based on OD600 and subsequently separated by sodium dodecyl sulfate polyacrylamide gel electrophoresis (SDS-PAGE) on 12% polyacrylamide gels at 120 volts in a Quadra Mini-Vertical PAGE/Blotting System (CBS Scientific). Resolved proteins were then transferred from polyacrylamide gels to 0.45 μm PVDF membranes (Immobilon, Roche) at 200 V for 1 hr. After transfer, membranes were stored overnight in TBST (0.1% Tween-20 (Amresco) in TBS (Corning)) at 4°C. Membranes were then blocked in TBST + 5% Bovine Serum Albumin (BSA) (Amresco) for 1 hour while rocking at room temperature. Rabbit anti-LukA (1:50000), rabbit anti-LukS-PV (1:50000), and rabbit anti-Hla (1:50000) antibodies (62) were incubated with the membranes in TBST for 1 hour. Although these antibodies recognize LukA, LukS-PV, and Hla they also permit detection of Protein A due to its binding of the FC region of antibodies (94). Following incubation with primary antibodies, membranes were washed three times in TBST for 15 minutes and goat anti-rabbit IgG (H+L) conjugated to Alkaline phosphatase (Thermo) was added at a 1:5000 dilution in TBST with 5% BSA for 1 hour. The membranes were washed 3 times in TBST for 15 minutes each and blots were developed with BCIP/NBT (5-bromo-4-chloro-3-indoyl-phosphate/nitro blue tetrazolium) color development substrate (VWR).

### AIP reporter assays

Reporter strains containing pDB59 (described above) and test strains (from which conditioned media was collected) were cultured overnight in TSB supplemented with Cm at 37°C with shaking at 200 rpm. The next day samples were diluted 1:100 in fresh TSB and incubated for 5 hours at 37°C with shaking at 200 rpm. Samples were centrifuged at 4200 rpm for 10 minutes. Supernatants from test strains were collected and filter sterilized through a 0.22 nm syringe filter and mixed 2:1 with fresh TSB. The reporter strain pellets were then resuspended in 5 mL of the 2:1 “conditioned medium:fresh medium” mix and incubated for an additional three hours. The bacteria were washed twice in 3.5 ml PBS and resuspended in 1mL of PBS. Samples were then diluted 1:1 in PBS and aliquoted in triplicate into a clear bottom 96-well black plate. Bacterial optical density (OD600) as well as GFP fluorescence (EX 495 nm, EM 509 nm) were measured using a Biotek Synergy H1 microplate reader. The data were normalized and displayed as GFP/OD.

### Generation of murine bone marrow derived macrophages

Murine bone marrow macrophages (BMM) were differentiated from the bone marrow of six to eight week old female or male C57BL/6J (WT), TLR2^−/−^, TLR4^−/−^, and MyD88^−/−^ mice as previously described (47).

### Macrophage cytokine Profiles

Bacteria were grown overnight in triplicate in a 96-well round bottom plate (Corning) in 150 μl of TSB at 37°C with shaking at 200 rpm. Samples were diluted 1:50 in 147 μl of fresh TSB in a 96-well round bottom plate and incubated for 8 hours at 37°C with shaking at 200 rpm. Bacterial growth was assessed by reading OD600 followed by centrifugation at 3700 rpm for 15 minutes to pellet cells. Supernatants were collected and stored at −80°C. Primary bone marrow derived macrophages (BMM) from WT, TLR2^−/−^, TLR4^−/−^, and MyD88^−/−^ mice were thawed and incubated at 37°C with 5% CO_2_ for 48 hours before seeding in a 96-well plate at 65000 cells per well. BMM were allowed to rest overnight before being treated with 10 μl of thawed bacterial supernatant for 24 hours at 37°C with 5% CO2. After 24 hours, 50 μl of supernatants were collected and stored at −80°C. To assess cytokine levels, a BDTM Cytometric Bead Array Mouse Soluble Protein Set and Soluble Protein Master Buffer Kit were used. The assay was conducted using the manufacturer’s instructions.

### Rabbit red blood cell lysis assay

For relative quantitation of hemolytic activity of WT *S. aureus* and mutants, strains were grown overnight in a round bottom 96-well plate in TSB with shaking at 37°C. The next day, overnights were diluted (1:50) in 150 µL TSB in a 96-well round bottom plate and incubated at 37°C with shaking for 6 hours. A 2% suspension of defibrinated rabbit red blood cells (RBCs) (Hemostat) was prepared by washing cells twice in 1x Phosphate Buffered Saline (PBS) and diluting in PBS to a final concentration of 2% RBCs from the stock RBC packed cell volume. This preparation was kept on ice until use. After 6 hours, the bacterial culture was centrifuged for 5 minutes at 3900 rpm. 100 µL of OD600 normalized cell-free supernatants were added to the top row of round bottom 96-well plates and a two-fold serial dilution series was set up with these supernatants. To each supernatant dilution, 2% RBCs were added (1:1) followed by incubation for 1 hour at 37°C. The plates were then centrifuged for 5 minutes at 1500 rpm to pellet non-lysed RBCs, supernatants were transferred to a flat bottom 96-well plate, and red blood cell lysis was measured at OD450 on a spectrophotometer.

### Murine skin and soft tissue infection model

Cultures of Wild type, Δ*mroQ*, Δ*mroQ+mroQ*, Δ*mroQ+mroQ(E141A)*, Δ*mroQ+mroQ(E142A)*, Δ*mroQ+mroQ(H180A)*, and Δ*agr* strains were inoculated from freshly isolated single colonies and grown with shaking at 200 rpm at 37°C overnight. The next day, a 1:100 subculture in 15 mL of TSB was incubated at 37°C for 3 hours with shaking. Cultures were then centrifuged for 5 minutes at maximum speed and cell pellets were washed twice in 5 mL PBS. Bacterial suspensions were diluted 2 mL into 8 mL of PBS and normalized with PBS to an OD600 of approximately 0.32-0.33 (1 × 10^8^ CFU/mL) and mixed 1:1 with sterile Cytodex beads (Sigma). Mice were deeply anesthetized with 2,2,2-tribromoethanol (Avertin; Sigma) via intraperitoneal injection followed by removal of fur and infecting with 100 μl PBS containing 1×10^7^ CFU of bacteria and Cytodex beads on the right and left side by intradermal injection. Mice were monitored daily; and at 72 or 120 hours post-infection, they were euthanized. Skin abscesses were collected, homogenized, spread onto TSA plates, and incubated overnight at 37°C in order to enumerate CFU. Skin homogenates were also used for cytometric bead array analysis of local cytokine levels as described above.

## Ethics Statement

All experiments were performed following the ethical standards of the institutional biosafety committee and the institutional animal care and use committee (IACUC) at Loyola University Chicago, Health Sciences Division. The institution is approved by Public Health Service (PHS) (#A3117-01 through 02/028/2022), is fully accredited by AAALAC International (#000180, certification dated 11/17/2016), and is registered/licensed by USDA (#33-R-0024 through 08/24/2020). All animal experiments were performed in ABSL2 facilities with IACUC approved protocols (IACUC #2017028) under the guidance of the office of laboratory animal welfare (OLAW) following the USDA and PHS Policy on Humane Care and Use of Laboratory Animals guidelines.

## Statistical Analyses

All experiments were repeated at least three independent times. For in vitro macrophage and AIP reporter data, statistical significance was analyzed from representative experiments conducted in triplicate and were repeated a minimum of three independent times. All statistical significance was analyzed using GraphPad Prism version 7.0 with statistical tests specified in the figure legends. Prior to conducting statistical analyses on data derived from animal studies, a D’Agostino and Pearson or Shapiro-Wilk normality test was conducted. Based on normality testing, either ANOVA or a non-parametric test was performed. Post-hoc statistical significance was calculated using Dunn’s post-hoc test. The number of animals per treatment group is indicated as “n” in the figure legends. For all other data, statistical significance (P < 0.05) was determined by one-way ANOVA with Tukey’s post hoc-test.

## Acknowledgments

We thank members of the Alonzo laboratory for helpful discussions and Alex Argianas for assistance with preliminary studies that helped guide project directions. This work was supported by grants NIH R01 AI120994 to FA, AHA 17PRE33660173 to JPG, and AHA 19POST34380259 to WPT.

